# FIVEx: an interactive multi-tissue eQTL browser

**DOI:** 10.1101/2021.01.22.426874

**Authors:** Alan Kwong, Andrew P. Boughton, Mukai Wang, Peter VandeHaar, Michael Boehnke, Gonçalo Abecasis, Hyun Min Kang

## Abstract

**Summary:** Expression quantitative trait loci (eQTLs) characterize the associations between genetic variation and gene expression to provide insights into tissue-specific gene regulation. Interactive visualization of tissue-specific eQTLs can facilitate our understanding of functional variants relevant to disease-related traits. However, combining the multi-dimensional nature of eQTLs into a concise and informative visualization is challenging. Existing eQTL visualization tools provide useful ways to summarize the unprecedented scale of transcriptomic data but are not necessarily tailored to answer questions about the functional interpretations of trait-associated variants or other variants of interest. We developed FIVEx, an interactive eQTL browser with an intuitive interface tailored to the functional interpretation of associated variants. It features the ability to navigate seamlessly between different data views while providing relevant tissue- and locus-specific information to offer users a better understanding of population-scale multi-tissue transcriptomic profiles. Our implementation of the FIVEx browser on the Gene-Tissue Expression (GTEx) dataset provides important insights for understanding potential tissue-specific regulatory mechanisms underlying trait-associated signals.

**Availability and implementation:** A FIVEx instance visualizing GTEx v8 data can be found at https://eqtl.pheweb.org. The FIVEx source code is open source under an MIT license at https://github.com/statgen/fivex.

## Introduction

Expression quantitative trait loci (eQTLs) are an important piece of the puzzle for understanding the regulatory mechanisms underlying genetic associations (Gallagher and Chen-Plotkin, 2018). Continuing advances in genomic technology have allowed researchers to generate enormous amounts of molecular profiles across many individuals and tissues. For example, the Genotype Tissue Expression (GTEx) Consortium analyzed transcriptomic profiles of 49 different tissues across 838 samples and identified >4 million eQTLs (Aguet, et al., 2019). The sheer number of eQTLs produced in such datasets require scalable, custom-designed visualization tools as aids for interpretation and analysis which will allow the exploration of a wide range of clinically relevant hypotheses, such as interpreting potential regulatory mechanisms in individual genome-wide association study (GWAS) signals (Roselli, et al., 2018; Yengo, et al., 2018), understanding tissue-specific epigenetic architecture of complex traits (Ehrlich, et al., 2019), and pinpointing likely causal variants by colocalizing GWAS and eQTL signals (Liu, et al., 2018; Wu, et al., 2019).

Interactive web applications such as the GTEx Portal (https://gtexportal.org) facilitate the functional interpretation of disease-associated regulatory variants. However, existing tools mainly focus on providing regional summaries of cis-eQTLs rather than tailored information relevant to functional interpretation. These tools also do not provide connections to other relevant online resources such as PheWeb (Gagliano Taliun, et al., 2020), BRAVO (http://bravo.sph.umich.edu), gnomAD (Karczewski, et al., 2020), and the UCSC browser (Haeussler, et al., 2019), making it challenging for users to gain a holistic understanding of the functional mechanisms underlying gene regulation.

To facilitate functional interpretation of regulatory variants from population-scale transcriptomic resources like GTEx, we developed FIVEx (Functional Interpretation and Visualization of Expression), an eQTL-focused web application that leverages the widely used tools LocusZoom (Pruim, et al., 2010) and LD server (https://github.com/statgen/LDServer). FIVEx visualizes the genomic landscape of cis-eQTLs across multiple tissues, focusing on a variant, gene, or genomic region. FIVEx is designed to aid the interpretation of the regulatory functions of genetic variants by providing answers to functionally relevant questions, for example, (1) how likely is a specific genetic variant to be causal for a cis-eQTL; (2) is a cis-eQTL tissue-specific or shared across tissues; (3) what is the linkage disequilibrium (LD) structure around the variant or gene; (4) which nearby genes are likely co-regulated by the variant and in which tissues; (5) whether there is additional information from other resources, such as biobank-based PheWAS results or regulatory genomic resources, that corroborates functional interpretation. FIVEx provides interactive visualizations of cis-eQTLs, capitalizing on functional interpretation and connecting to relevant external resources (Figure S1–15). We expect FIVEx to serve as a useful community resource of multi-tissue eQTLs, complementing the GTEx Portal, the OpenTargets Platform (Carvalho-Silva, et al., 2019), and other widely used web tools.

### Key Features

The primary goal of FIVEx’s visualizations is to aid exploratory analysis to help investigators interpret the regulatory functions of the variants identified from GWAS or other genetic studies (Figure S1). FIVEx allows investigators to query an eQTL database for a variant, gene, or region to interactively visualize multi-tissue cis-eQTLs from various viewpoints, either in a single-variant view of all cis-eQTLs, or in multiple LocusZoom cis-eQTL plots for selected genes and tissues. When a variant is queried, FIVEx visualizes the landscape of cis-eQTLs associated with the variant across all nearby genes and all tissues. FIVEx also highlights the variants most likely to have causal regulatory effects. When a gene or a region is queried, FIVEx offers multi-tissue and/or multi-gene LocusZoom visualizations of strongly associated cis-eQTLs, along with a summary list of variants that most likely regulate genes across the tissues. By effectively visualizing GTEx data, FIVEx can help unravel potential tissue-specific regulatory effects underlying the association (Figure S2–S4).

FIVEx’s variant-centric visualization helps users interpret the potentially causal role of a variant in tissue-specific regulation. For example, if a user queries for rs934197, a strong known association signal for low density lipoprotein (LDL) in *APOB*, FIVEx provides a comprehensive view of all cis-eQTLs of nearby genes, including *APOB* across multiple tissues (Figure S12). FIVEx provides posterior inclusion probabilities (PIPs) (Wen, et al., 2017) to inform whether the variant is likely a causal variant. For example, rs934197 shows the strongest p-value (4×10^−17^) in subcutaneous adipose but shows low PIP (<.001), suggesting that the association is likely a shadow signal of a stronger eQTL. However, in other tissues, such as tibial artery and esophagus, rs934197 has highest PIP despite weaker marginal p-values indicating a potentially causal regulatory role for *APOB* in those tissues.

FIVEx’s region view can help us explore the cis-eQTL landscape around the variant of interest in more detail. Users can interactively add multiple panels of eQTL LocusZoom plots, for example, one per tissue for a given gene (Figure 1A). In the *APOB* example, FIVEx helps visualize that several tissues in gastrointestinal and lower circulatory systems share the same peak cis-eQTL at rs934197, suggesting colocalization between *APOB* regulation and blood cholesterol associations in these tissues.

**Figure 1.**
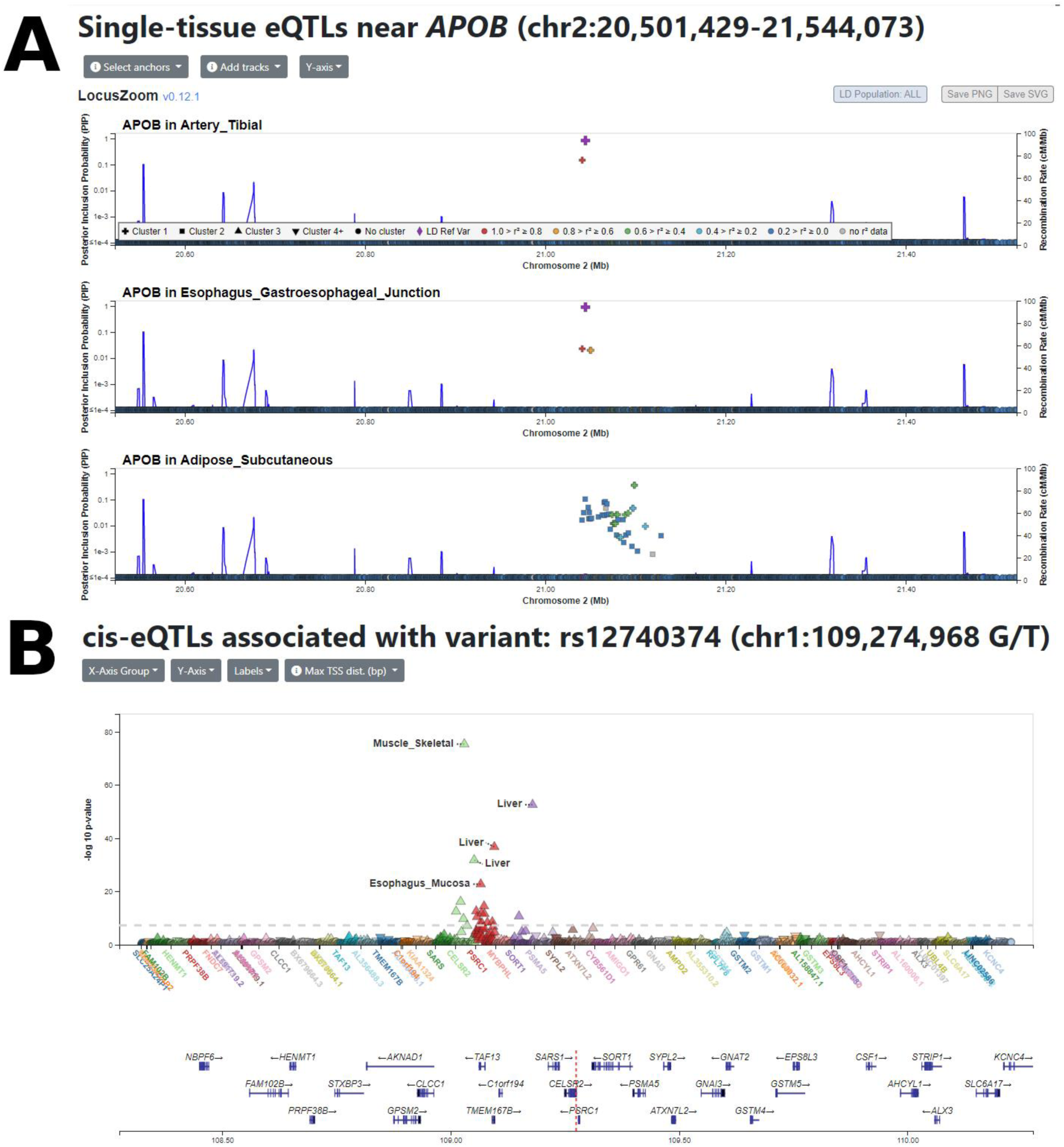
Examples of FIVEx views. (A) A locus-centric view of FIVEx when querying *APOB* after adding additional tracks with relevant tissues, showing cis-eQTLs for different tissues near *APOB*. The first two tissues share the same peak cis-eQTLs, while the other tissue does not. The points’ colors indicate LD with respect to the index variant, while the points’ shapes indicate PIP cluster membership. Each cluster represents an association, with the strength of the PIPs reflecting each point’s contribution towards that association. (B) Another variant-centric view of FIVEx when querying rs12740374, the most strongly associated variant with self-reported high cholesterol level in UK Biobank near the *SORT1* locus. This variant is a strong cis-eQTL regulating multiple genes (*SORT1, PSRC1, CELSR2*), particularly in liver tissue, illustrating the benefit of using FIVEx to identify co-regulation of proximal genes.

In addition to visualizing multi-tissue cis-eQTLs for the same gene, FIVEx can also show co-regulation of multiple nearby genes in the same tissue. For example, rs12740374 shows the strongest association with high cholesterol levels in the UK Biobank near the *SORT1-PSRC1-CELSR2* locus, and previous studies suggest liver-specific regulation as a possible functional mechanism (Musunuru, et al., 2010; Schadt, et al., 2008; Wang, et al., 2018). FIVEx’s single variant view clearly illustrates liver-specific cis-eQTLs in these genes for rs12740347 with strong PIPs (Figure 1B). The region view clearly illustrates that the variant is the top signal for *SORT1* and *PSRC1*, and the second top signal for *CELSR2* (Figure S15). Through interactive eQTL navigation, FIVEx provides additional context for interpreting association signals to guide hypotheses for underlying regulatory mechanisms.

A more detailed description of these example usages can be found in the Supplementary Text and in an online tutorial at https://eqtl.pheweb.org/tutorial.

## Discussion

Understanding the function of trait-associated non-coding variants is becoming increasingly important as more genomes, transcriptomes, and epigenomes are sequenced. Gene regulation is believed to be involved in a large fraction of such associations, but there are limited resources which investigators can use to generate hypotheses for explaining regulatory mechanisms underlying association signals. FIVEx offers new interactive ways to visualize and summarize eQTLs in a tissue-specific manner by combining key features from LocusZoom and PheWeb, focusing on putative causal eQTLs through PIPs (Figure S4). We expect FIVEx will aid with translating GWAS associations into underlying regulatory mechanisms by enabling the exploration of plausible hypotheses through our intuitive and practical user interface. As more online resources like FIVEx become available to address tailored scientific questions on functional variants, we expect that precise and integrative translation of genomic findings will be more accessible to the broader scientific community.

## Supplementary Text

### Overview

In this supplementary text, we elaborate on the examples of using FIVEx described in the main text with more detail to clarify the functionalities provided by our web app. Most of these examples are also illustrated in our online video tutorial at https://eqtl.pheweb.org/tutorial

### Interpreting LDL cholesterol association near *APOB*

To illustrate the capabilities of FIVEx, we use low-density lipoprotein (LDL) cholesterol level in blood, a trait strongly associated with cardiovascular disease (Klarin, et al., 2018), as a motivating example. In a UK Biobank (UKB) GWAS of LDL cholesterol levels (NRM_LDL_C), one of the strongest association signals is located near *APOB* with the minimum p-value 5.1×10^−27^ at rs934197 (Figure S1A-B). Due to the strength of the association signal and the long-range LD in this region, it is unclear which variant is causal and what the underlying mechanism is behind the *APOB* association (Niu, et al., 2017; Sabatti, et al., 2009; Willer, et al., 2008).

If a user searches for rs934197, the peak SNP for *APOB*, FIVEx visualizes a landscape of cis-eQTL associations with the variant across nearby (<1Mb) genes and tissues (Figure 1A, S5–6). Interestingly, rs934197 shows strong associations with *APOB* expression levels in several tissues, including subcutaneous adipose (p = 4.0 × 10^−17^), tibial artery (p = 7.9 × 10^−11^), and esophagus gastroesophageal junction (p = 6.9 × 10^−9^), but this does not automatically mean that the variant directly regulates *APOB* expression levels. Instead, the associations may be explained as indirect shadows of other causal variants via LD. However, existing eQTL browsers do not offer straightforward ways to distinguish putatively causal associations from indirect signals. In contrast, FIVEx allows users to distinguish between the two by providing posterior inclusion probabilities (PIPs) under the DAP-G model (Wen, et al., 2017). In the *APOB* example, FIVEx illustrates that the strongest eQTL found in subcutaneous adipose is unlikely to be a regulatory variant (PIP ≈ 0) (Figure S7), but that instead the eQTL is better explained as a shadow signal of another cis-eQTL, most likely rs4665178 (p = 1.4×10^−28^, PIP = 0.35), located 54kb upstream. Meanwhile, rs934197 has the strongest PIPs in esophagus gastroesophageal junction (0.90), esophagus muscularis (0.71), tibial artery (0.83), and sigmoid colon (0.73) tissues, even though their marginal p-values were weaker than the p-value for subcutaneous adipose. These results illustrate how FIVEx can help characterize tissue-specific cis-regulation potentially caused by a specific variant more comprehensively than existing tools (Figure S8–10).

### Multi-tissue visualization of tissue-specific regulation of *APOB*

When a gene is queried, FIVEx displays multiple parallel LocusZoom region plots to visualize cis-eQTLs across tissues and/or nearby genes to visualize the complex structure of gene regulation entangled with LD, along with the list of most likely causal variants (i.e. those with highest PIPs) (Figure S11). Users can interactively add tissues or nearby genes to the LocusZoom plot (Figure 1B, S12), or change the index variant to visualize whether it aligns with the peak cis-eQTLs in each plot (Figure S13). In the *APOB* example, rs934197 is clearly the top signal for four tissues in the gastrointestinal or lower circulatory systems, while other tissues—heart (rs661665), adipose (rs4665178), skin (rs579826), and muscle (rs56327713)—feature different nearby top variants (Table S1). Among these variants, only rs934197 co-localized as a peak genome-wide significant association signal with any phenotype in UKB PheWAS, which can be easily seen via the link FIVEx provides to those results. Together, these observations suggest that the lipid association signal near *APOB* may be explained by gene regulation in specific gastrointestinal and/or circulatory systems.

### Tissue-specific co-regulation of multiple genes in *SORT1-PSRC1-CELSR2* locus

FIVEx also allows users to identify and visualize nearby genes sharing a cis-eQTL. For example, one of the peak signals associated with self-reported high cholesterol in the UK Biobank (biobank_20002_1473) is rs12740374 near the *SORT1-PSRC1-CELSR2* locus (Figure S3A). Querying this variant in FIVEx demonstrates that the variant is also a strong cis-eQTL, specifically in liver (Figure S14–15). Moreover, this variant is a likely causal regulatory variant for all three genes (*SORT1, PSRC1, CELSR2*) in liver, with very strong PIPs (≥0.99) for *SORT1* and *PSRC1*, and moderate PIP (0.42) for *CELSR2* (Figure S15). The cis-eQTL appears to be highly liver-specific for *SORT1* and *PSRC1*, consistent with the differential transcriptional activity observed in human hepatoma cells for both genes (Musunuru, et al., 2010; Schadt, et al., 2008; Wang, et al., 2018). In contrast, for *CELSR2*, rs12740374 has modest (0.21 – 0.56) PIPs across 5 tissues, including liver (Figure S3B-D). This observation may serve as an important guide to generate hypotheses about the functional mechanisms regulating these lipid-associated genes. As shown by these examples, FIVEx provides investigators intuitive and interactive means to navigate eQTL data focusing on questions directly relevant to gene regulation. FIVEx serves as a complementary tool to the GTEx Portal and other online resources specifically in pinpointing putative functional variants through gene regulation among numerous trait-associated variants.

### Summary of features offered by FIVEx

FIVEx’s single variant view, which will be used when a user searches for a variant by position or rsID (Figure S5), provides the following functionality:

- Show eQTLs across all tissues for genes within 1MB of the variant (Figure 1B)
- Dynamically change the grouping of eQTLs by tissue, biological system, or gene (Figure S6)
- Dynamically switch between displaying p-values, effect sizes, and posterior inclusion probabilities (PIPs) in the same plot (Figure S7)
- Show or hide labels for top eQTLs (Figure S8)
- Change the window size around the current variant for eQTLs to display (Figure S9)
- Highlight eQTLs which affect the same gene, or those found in the same tissue (Figure S10)

FIVEx’s region view, which will be used when a user searches for a gene or a region, provides the following functionality:

- View a genomic region (locus plot) for the gene and tissue with the strongest association signal, along with a table of the most likely causal variants according to PIP (Figure S11)
- Add additional gene or tissue data tracks (Figure 1A, S12)
- Change the index variant for showing linkage disequilibrium in the region (Figure S13)
- Dynamically switch the displayed Y-axis variable for all tracks (Figure S14–15)

## List of Supplementary Figures

**Supplementary Figure S1.**
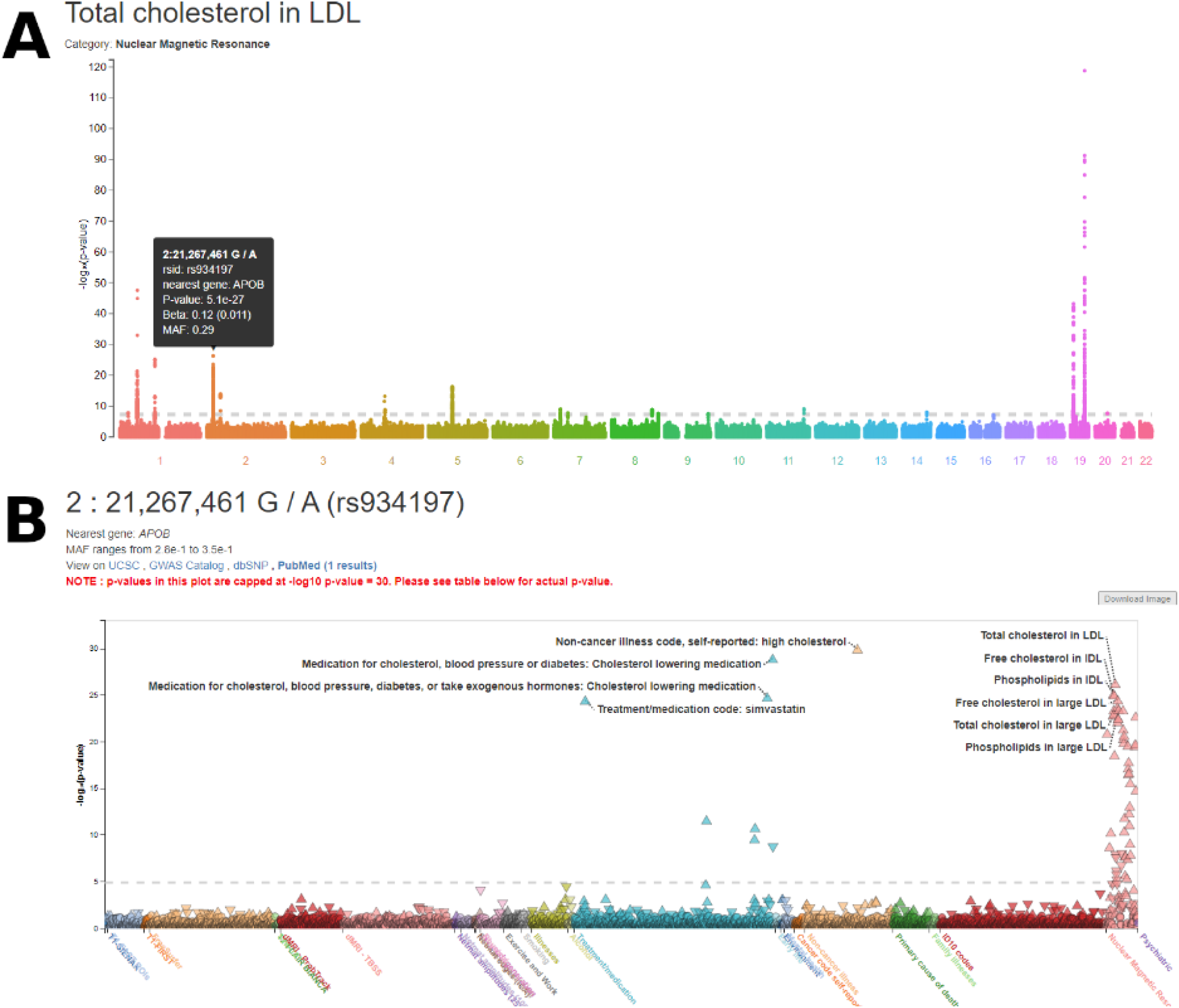
Exploring LDL-associated eQTLs in the *APOB* locus. (A) Using an external resource (the Oxford Brain Imaging Genetics Server) to search for variants associated with total cholesterol in LDL, the variant with the strongest association on chromosome 2 is rs934197, in the *APOB* locus. (B) A PheWAS view of this variant reveals multiple strong association signals, with both traits and medication use directly related to LDL cholesterol.

**Supplementary Figure S2.**
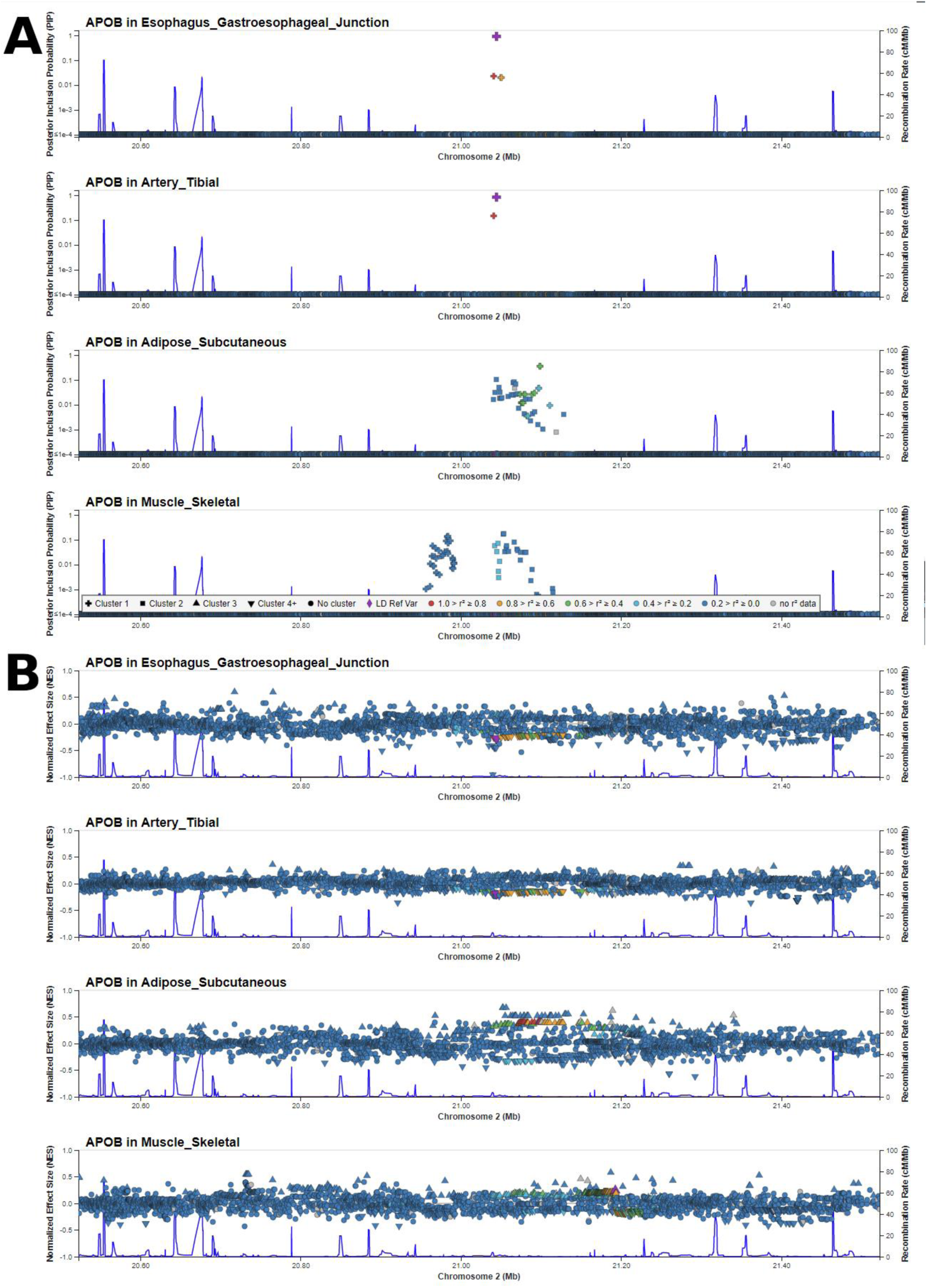
FIVEx provides genomic context for tissue-specific expression. (A) We view PIPs for different tissues in the *APOB* locus in FIVEx’s region view and use the top signal in esophagus – gastroesophageal junction (EGJ) tissue as the reference variant to compare with signals in other tissues. We see that while this is also the top variant in tibial arterial tissue, it is distinct from the signal cluster in subcutaneous adipose and skeletal muscle, showing tissue-specific expression differences associated with distinct LD blocks. (B) When we view by effect sizes, we see that variants associated with lower *APOB* expression in EGJ and tibial artery tissues, but higher expression in subcutaneous adipose tissue, highlighting the heterogeneous effects one variant can have on the same gene in different tissues.

**Supplementary Figure S3.**
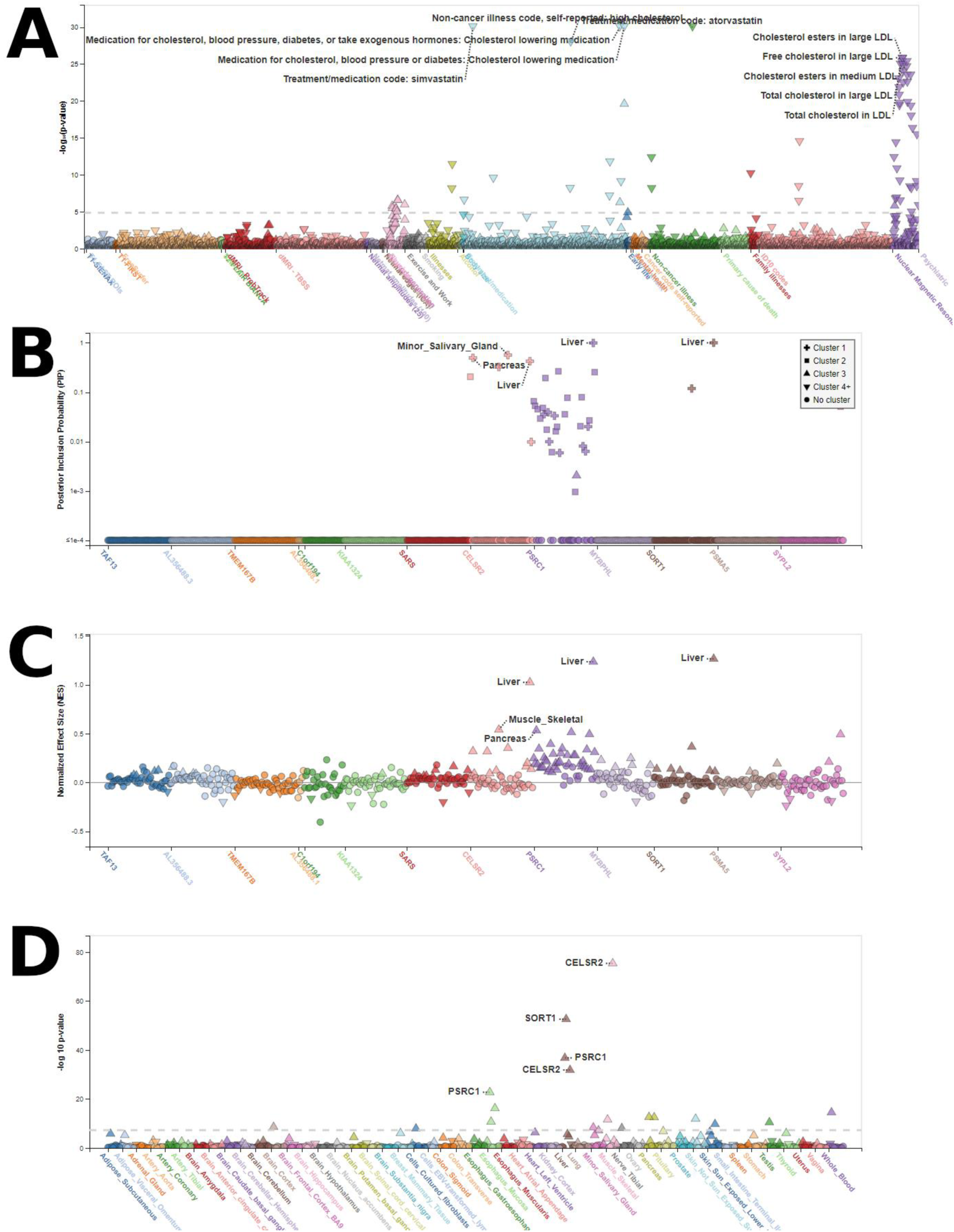
FIVEx shows multiple genes associated with a single variant. (A) Using UK Biobank as a reference, we searched for variants strongly associated with cholesterol-related traits. The variant rs12740374 in the *SORT1-PSRC1-CELSR2* locus has strong associations with multiple cholesterol-related traits. (B) In FIVEx’s single variant view with PIPs on the y-axis, we can see strong associations with liver tissue for all three genes, along with multiple other tissues, evidence for both tissue-specific and cross-tissue regulatory effects. (C) The pattern of effect sizes of the associations indicate that this variant has strong upregulation effects on all three genes in liver tissue. (D) The strong P-values in liver across all three tissues provide additional evidence for liver-specific regulatory mechanisms with this variant.

**Supplementary Figure S4.**
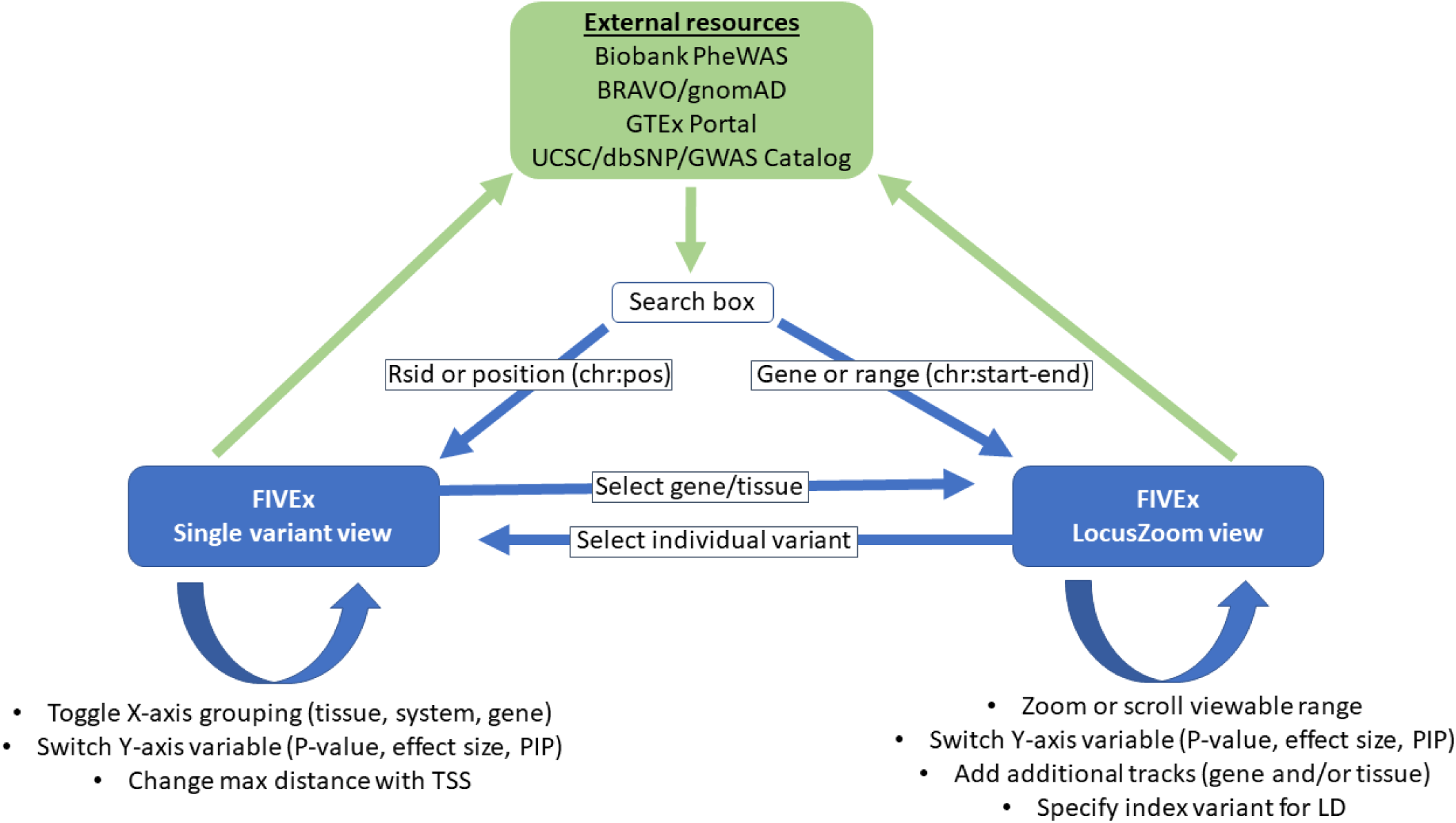
Navigating FIVEx. An illustration of navigating eQTL data in FIVEx. A researcher may wish to learn more about gene regulation related to a variant, gene, or region of interest. A variant query will send the user to a single variant view with extensive information about all eQTLs related to one variant, while a gene or region query will send the user to a LocusZoom view of the gene or region for the tissue with the strongest eQTL association. The user can manipulate the displayed data within each view in real time to show different metrics of association: P-values for evidence of association, effect sizes for strength of regulatory effect, or PIPs for a modeled probability of a variant being causal for an eQTL.

**Supplementary Figure S5.**
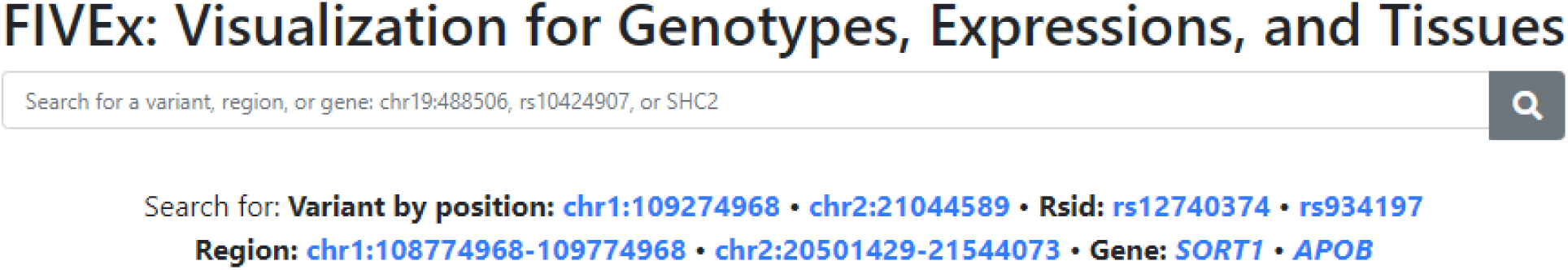
Step-by-step tutorial on reproducing examples in the FIVEx main text. The FIVEx homepage includes a search box which can find variants using chromosomal position or rs numbers. We first enter ‘rs934197’, the rsID for the top LDL-associated variant located ∼500 bp upstream of *APOB*, in the search box. FIVEx sends us to a single variant view.

**Supplementary Figure S6.**
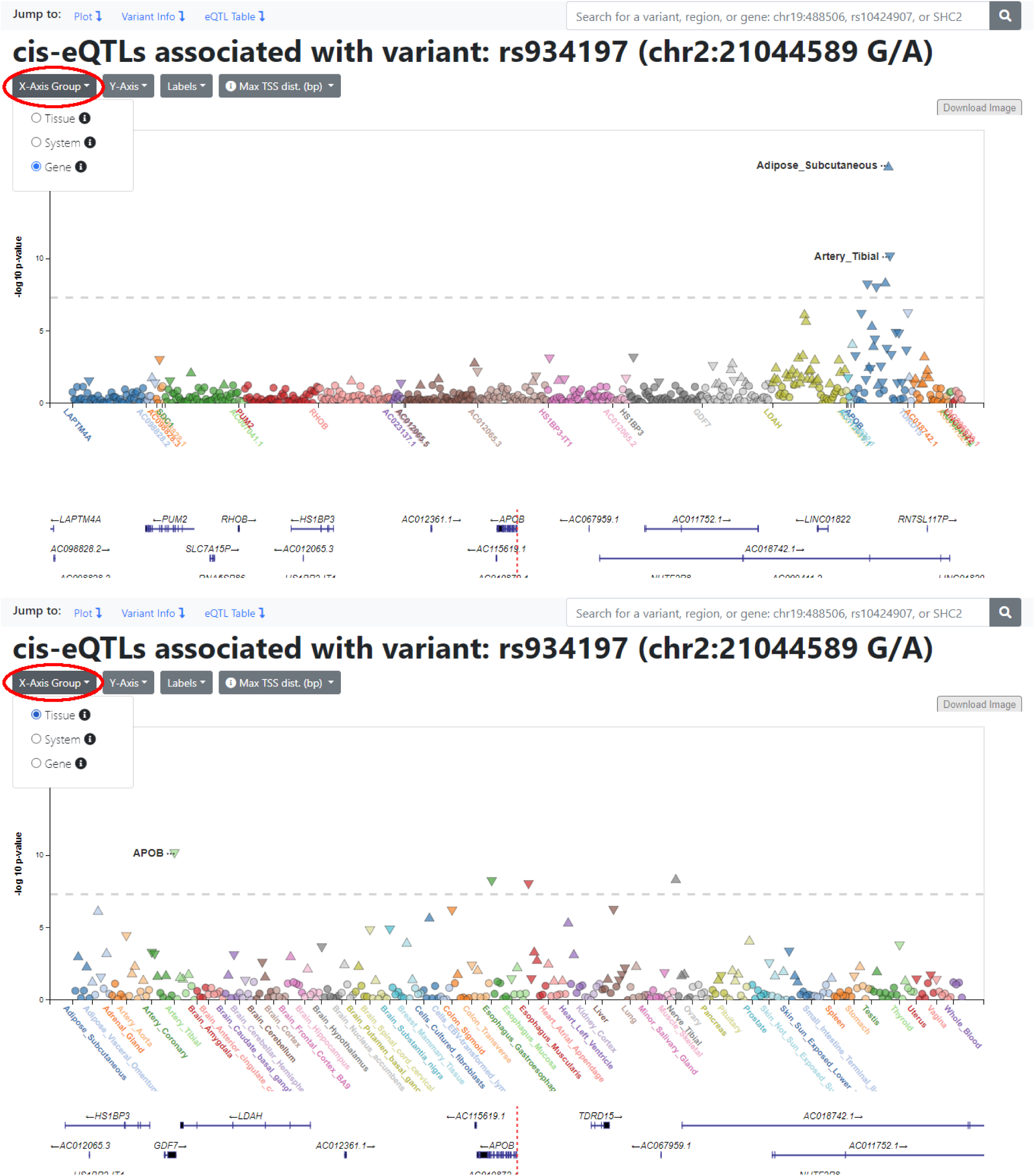
Dynamically change the grouping variable on the x-axis. The first dropdown menu allows us to change the x-axis grouping of the data in real time. Currently, the points are grouped by gene, arranged by the genomic positions of their TSS. Grouping by gene makes it easier to see multi-tissue regulation of specific genes, while grouping by tissue makes it easier to see multi-gene regulation in specific tissues. Grouping by system further groups tissues for a more general overview of gene regulation in different tissue systems.

**Supplementary Figure S7.**
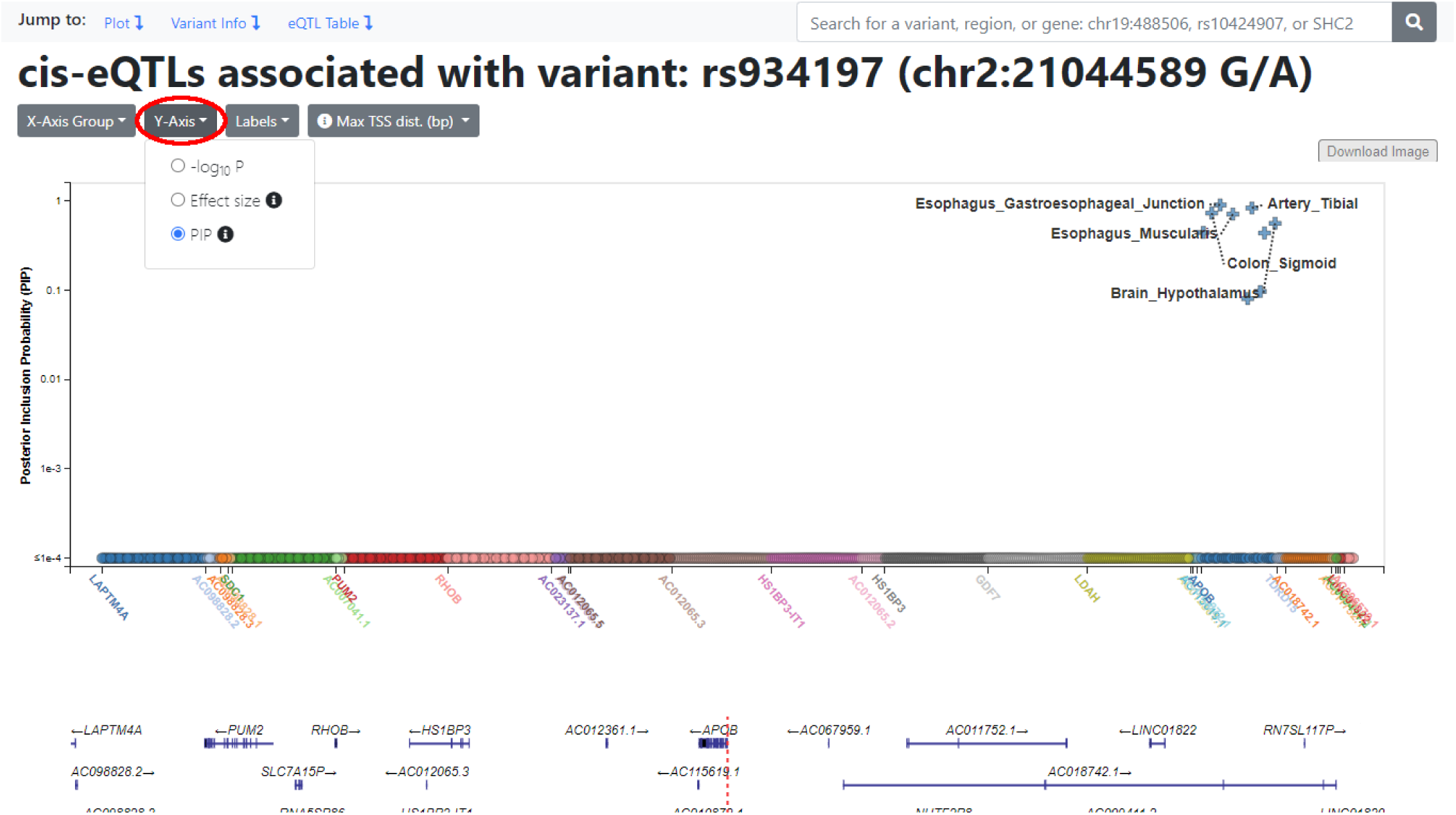
Dynamically changing the displayed Y-axis variable. The second dropdown menu allows us to show y-axis variables other than P-values. For example, we can view PIPs, which take into account the LD structure around a variant to calculate a posterior probability that a variant is causal for an eQTL. For example, subcutaneous adipose, which has a highly significant P-value for its association with the expression of *APOB*, does not have a corresponding PIP signal.

**Supplemental Figure S8.**
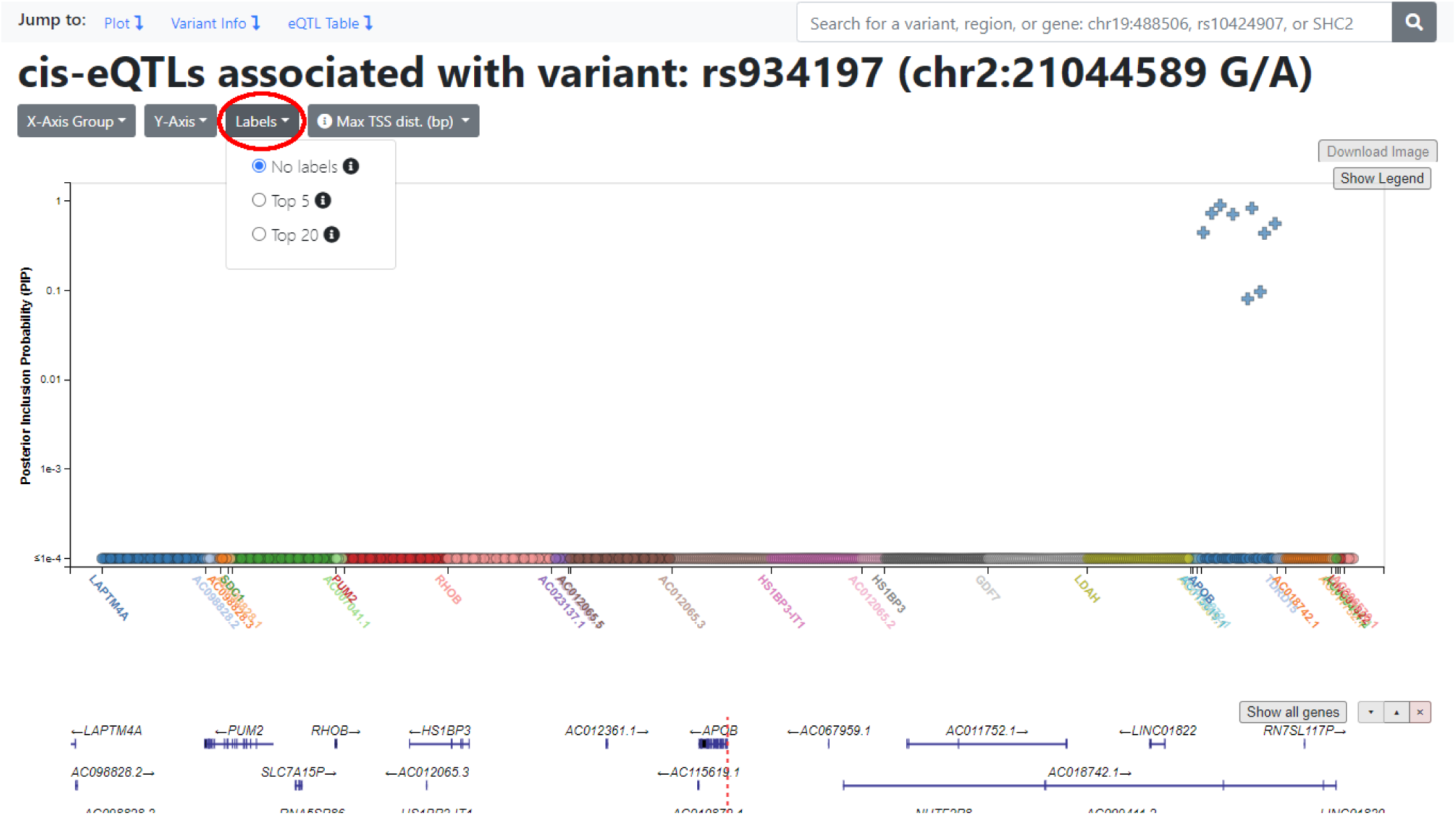
Toggling labels for the strongest signals. The third dropdown menu allows us to toggle labels for significant signals. Here we turned off the labels to give us a better look at the PIP signals in *APOB*. When the results are grouped on the x-axis by gene, labels show the tissues for the data points; when the results are grouped by tissue or system, labels show genes instead.

**Supplementary Figure S9.**
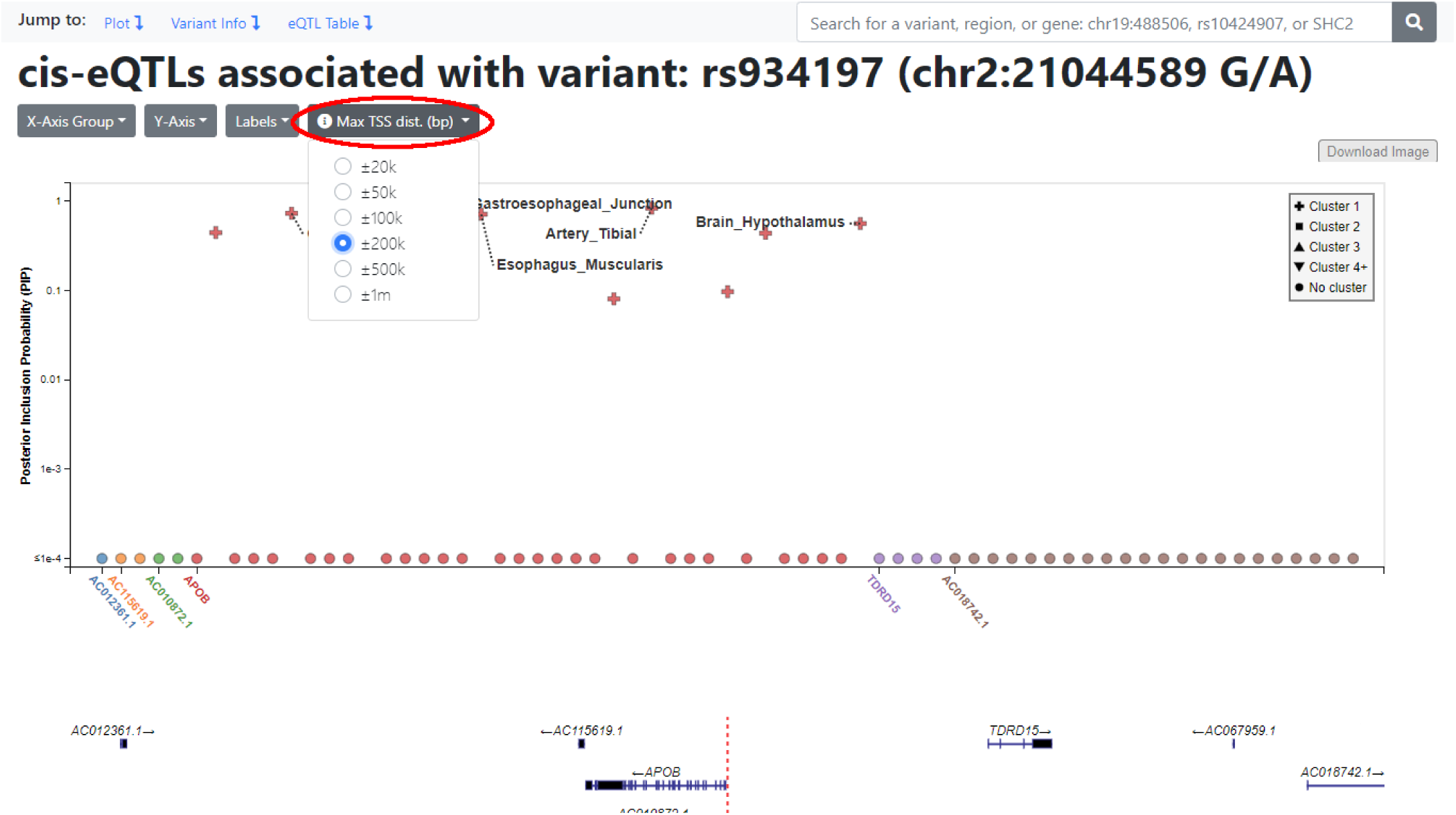
Changing the viewing window using maximum TSS distance. The fourth dropdown menu allows us to change the amount of data displayed by setting a maximum transcription start site (TSS) distance from the current variant. For example, setting this to ±200k means displaying only the eQTLs for genes with a TSS within 200kbp of the current variant.

**Supplementary Figure S10.**
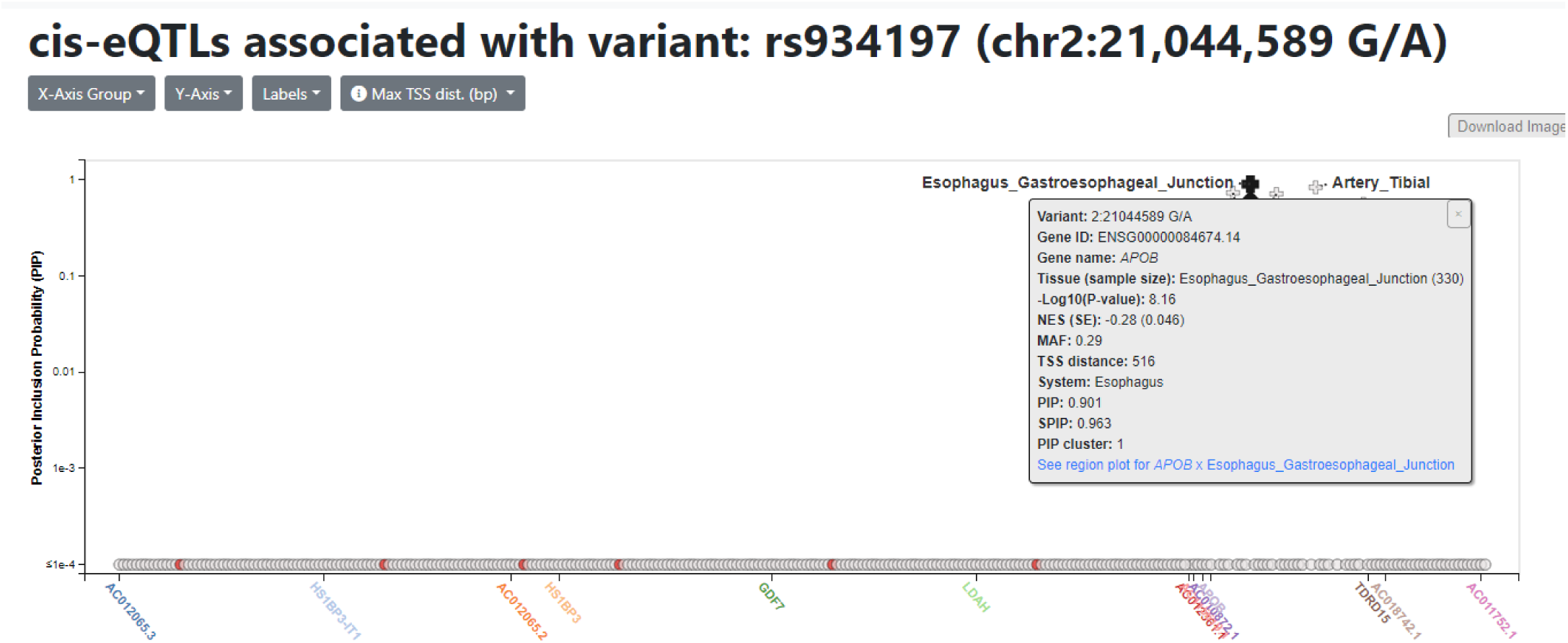
Dynamic highlighting of points with a shared attribute. Clicking on any point highlights all other points which share an attribute which is not the current X-axis grouping variable, allowing us to easily spot data points of interest. For example, if we are using tissue as the X-axis grouping variable, then clicking on a point will highlight all points which affect the same gene. Here, since we are using genes as our X-axis grouping variable, when we click on an eQTL which affects gene expression in subcutaneous adipose tissue, we see all other eQTLs in subcutaneous adipose tissue highlighted in red. Clicking on any point also brings up a tooltip window displaying detailed information about that eQTL, with a link to a region view around this signal. We will follow the link to a LocusZoom view of the region around rs934197 in gastroesophageal junction tissue.

**Supplemental Figure S11.**
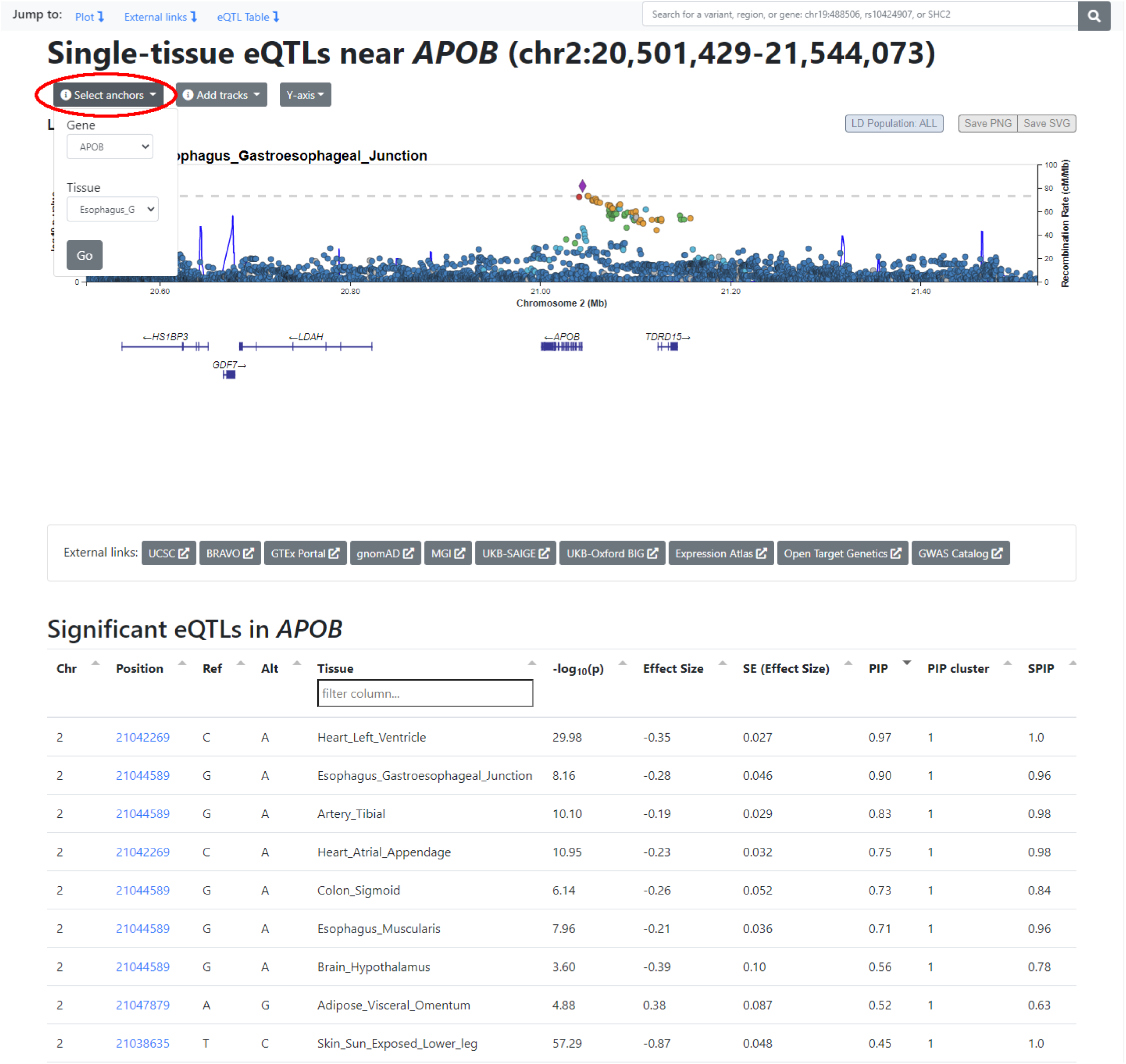
A LocusZoom region view is defined by an anchor gene and tissue. The LocusZoom view around our variant is “anchored” around one gene and one tissue. In our example, the anchors are *APOB* and Esophagus – Gastroesophageal Junction, respectively. Anchors give us stable pivot points around which we can explore additional genes and tissues. We can change our anchor gene and tissue via the first pulldown menu. A table of PIP signals for the anchor gene is shown below the region plot, sorted by decreasing PIP signal strength.

**Supplementary Figure S12.**
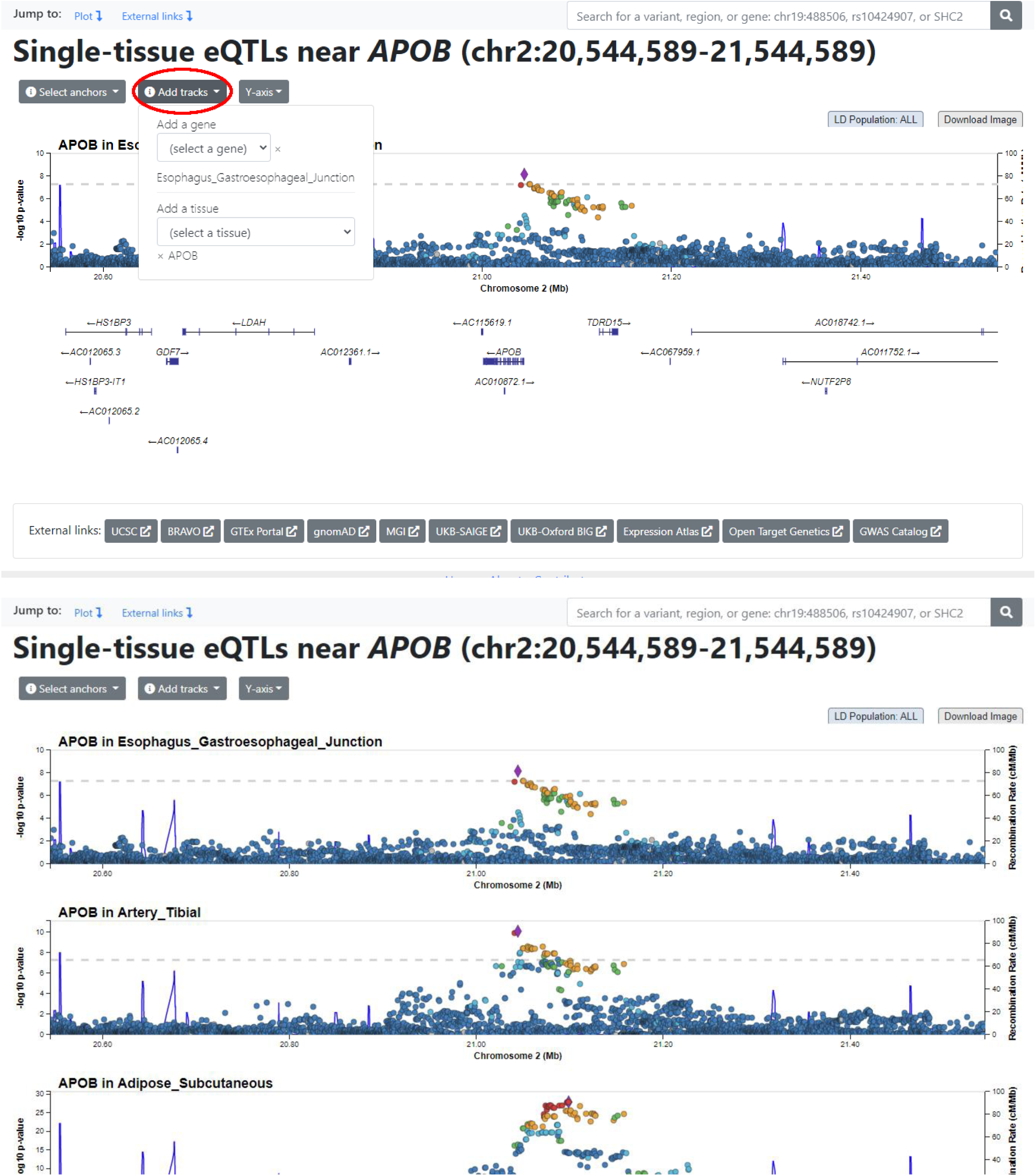
Adding additional tracks in LocusZoom view to facilitate comparison. The second pulldown menu allows us to add genes or tissues with respect to one of the anchors. For example, we can add a track to see eQTLs for *APOB*, our anchor gene, in a different tissue. Similarly, we can add a track to see eQTLs for a different gene in our anchor tissue. We will add tibial artery and subcutaneous adipose as additional tissue tracks.

**Supplementary Figure S13.**
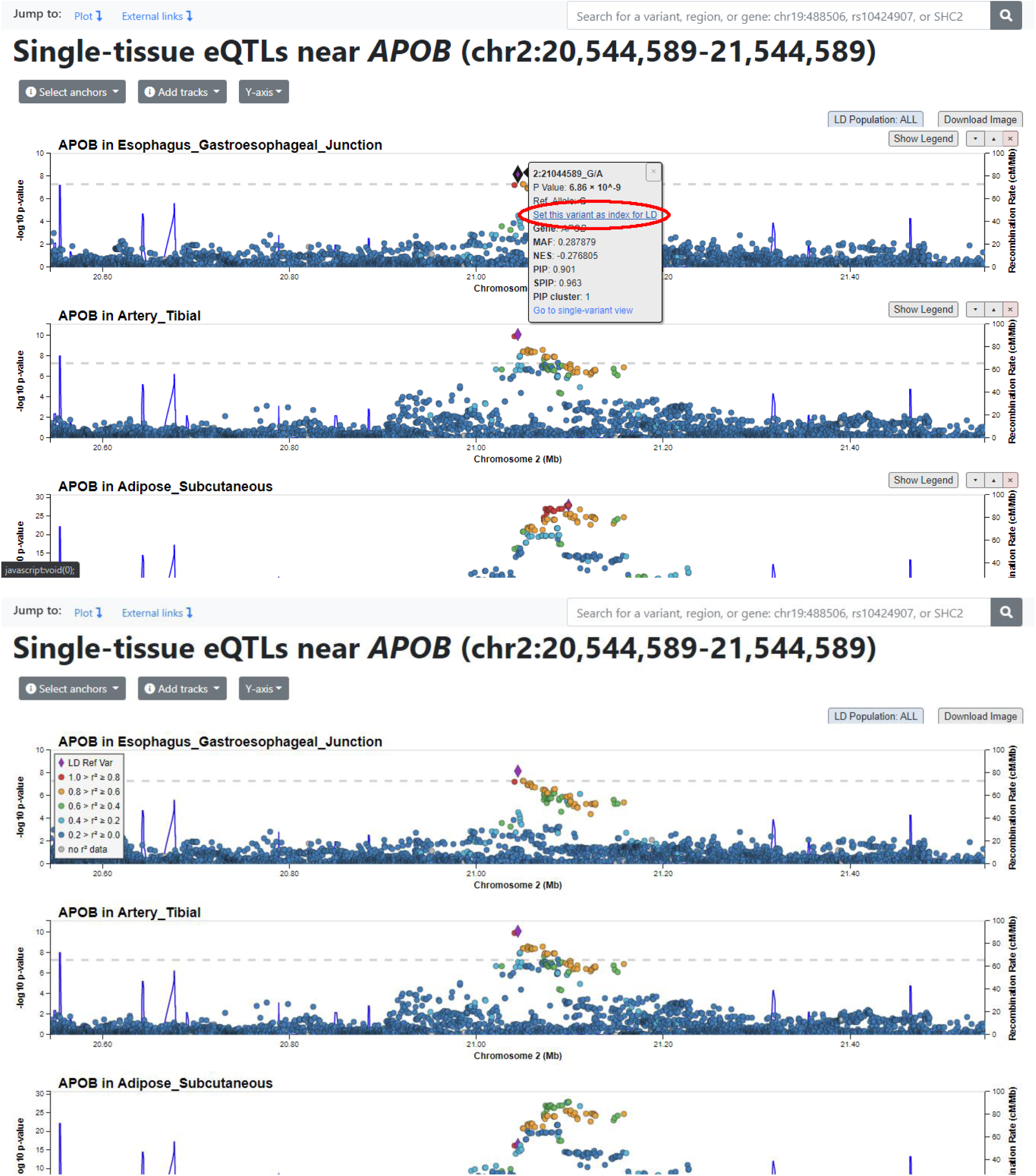
Setting a reference variant to obtain linkage disequilibrium information. The tooltip window in region view allows us to set a variant as the index for linkage disequilibrium (LD), which recolors the points in all the displayed plots to reflect their LD with the index. LD information is provided by the LD server and is currently based on data from the 1000 Genomes Project. The index variant is indicated by a purple diamond. We see that our index variant is the top signal in esophageal and arterial tissues but is a shadow of a stronger signal in adipose.

**Supplementary Figure S14.**
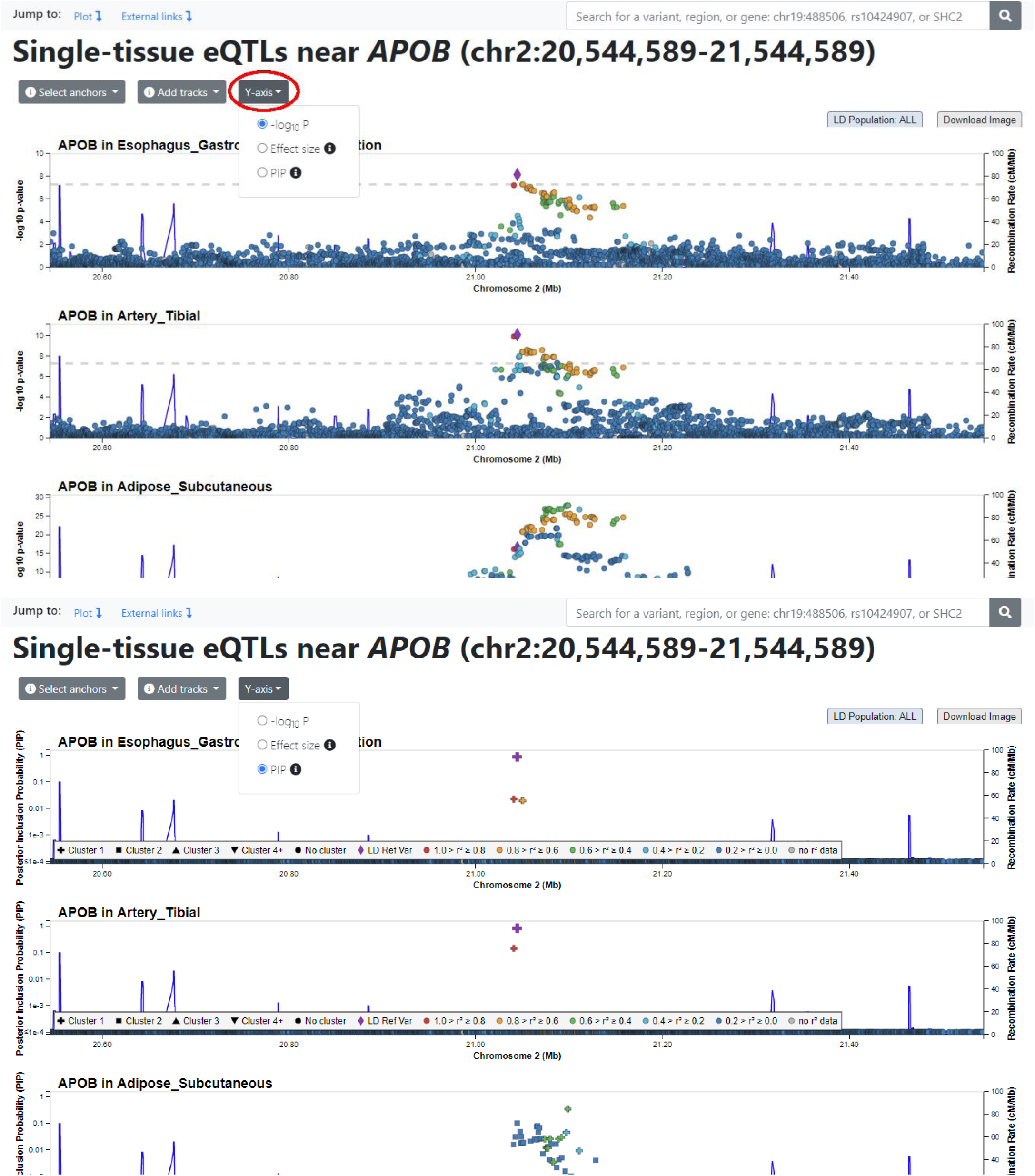
Dynamic y-axis variables in LocusZoom view. The third pulldown menu changes the Y-axis variable just like in our single variant view. In PIP view, we see that in esophageal and arterial tissues, there is only one signal cluster and our index variant is the strongest contributor. However, in adipose, the PIP signal is distributed more evenly between two different variant clusters, indicating more uncertainty about the potentially causal variant for eQTLs in this tissue.

**Supplementary Figure S15.**
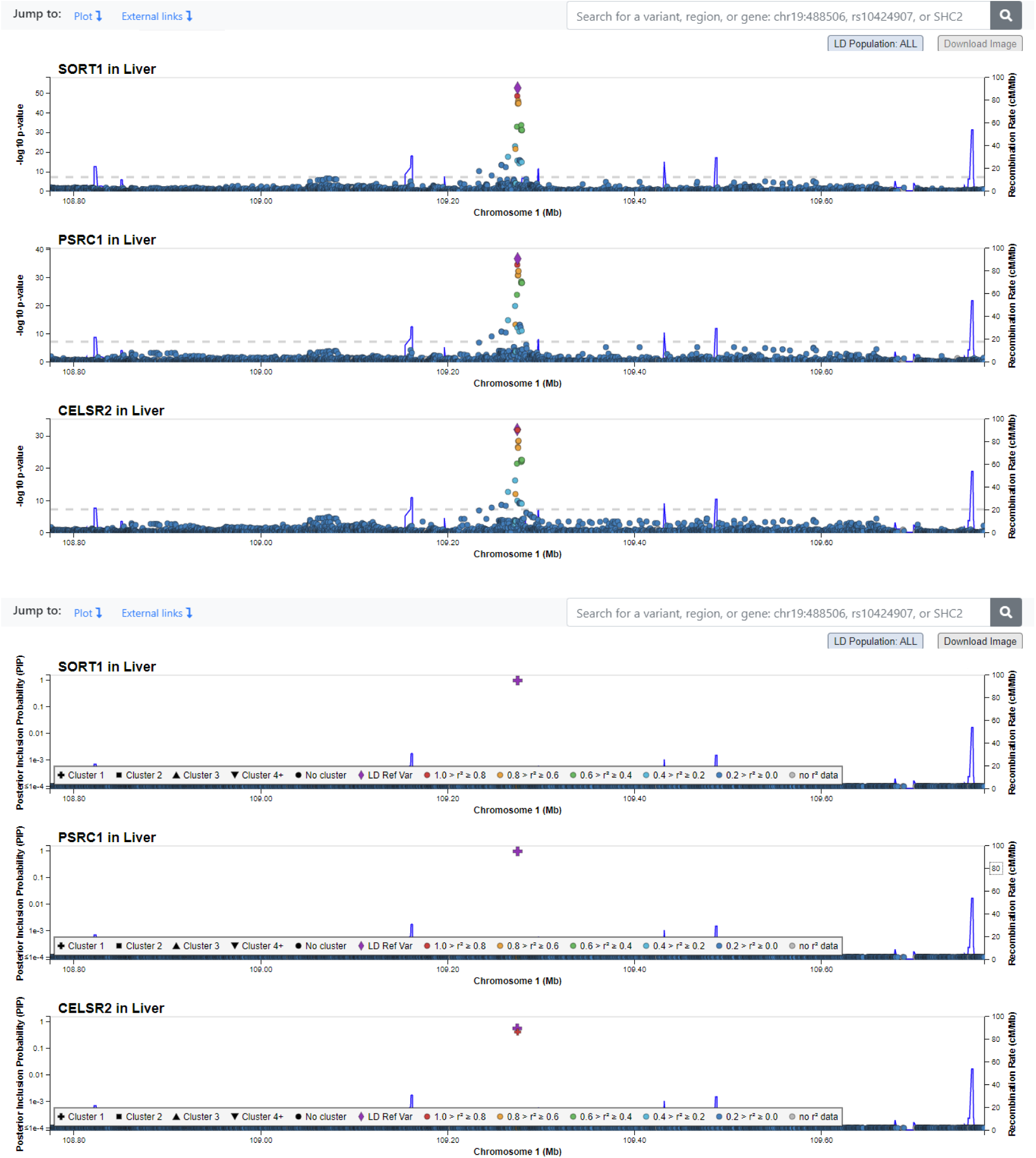
Comparing expressions of multiple genes in one tissue via LocusZoom view. We can also compare the eQTL signals for multiple genes in the same tissue. Here, we searched for the top variant for cholesterol-related association signals in the *SORT1-PSRC1-CELSR2* locus, rs12740374, and navigated to a region view. Using liver tissue and *SORT1* as anchors, we added the other two genes to compare the eQTL signals between them. We see that rs12740374 is the strongest signal for these three genes, both for P-values and PIPs, suggesting a regulatory pathway involving all three.

**Supplementary Table S1.**
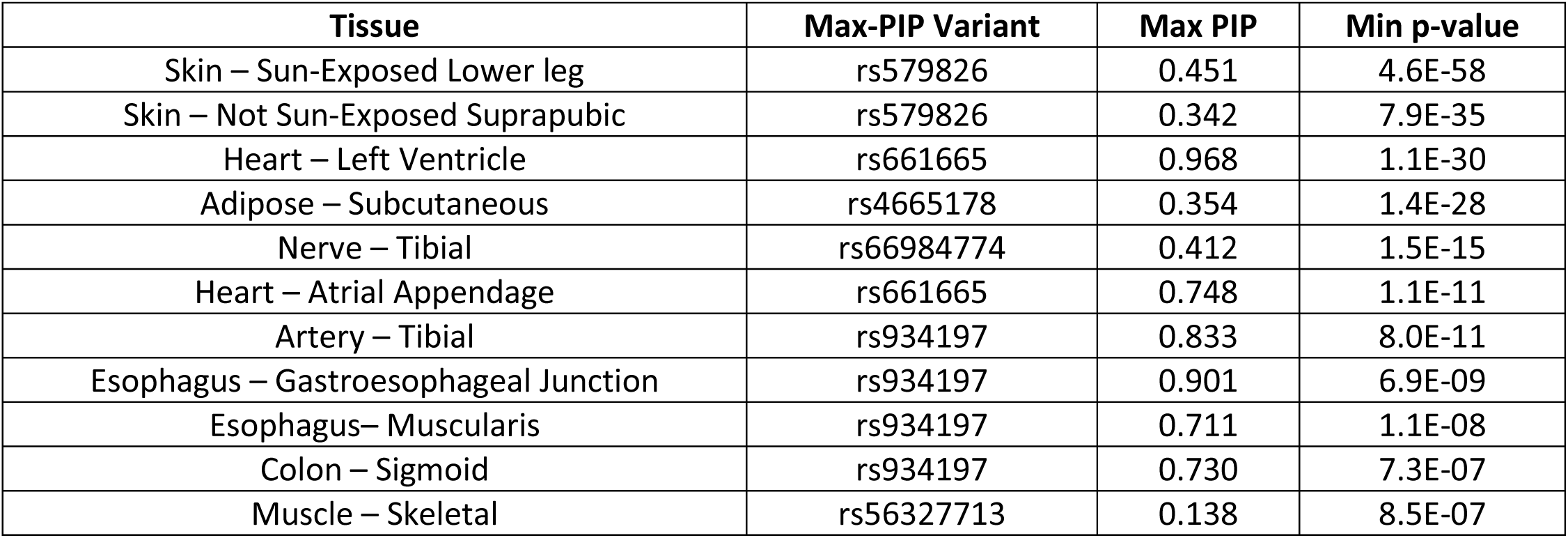
Comparison of top PIP variants for *APOB* across different tissues (p < 10^−6^). Tibial artery, gastroesophageal junction, sigmoid colon, and skeletal muscle tissues share the same top PIP signal (rs934197) for affecting the expression of *APOB*. Both skin tissues share a different variant as the top signal (rs579826), while subcutaneous adipose tissue has a third variant (rs4665178) with the strongest PIP. This highlights differences in regulatory mechanisms between different tissues for the same gene, providing clues for a better understanding of tissue-specific gene expression.

| **Tissue** | **Max-PIP Variant** | **Max PIP** | **Min p-value** |
| --- | --- | --- | --- |
| Skin – Sun-Exposed Lower leg | rs579826 | 0.451 | 4.6E-58 |
| Skin – Not Sun-Exposed Suprapubic | rs579826 | 0.342 | 7.9E-35 |
| Heart – Left Ventricle | rs661665 | 0.968 | 1.1E-30 |
| Adipose – Subcutaneous | rs4665178 | 0.354 | 1.4E-28 |
| Nerve – Tibial | rs66984774 | 0.412 | 1.5E-15 |
| Heart – Atrial Appendage | rs661665 | 0.748 | 1.1E-11 |
| Artery – Tibial | rs934197 | 0.833 | 8.0E-11 |
| Esophagus – Gastroesophageal Junction | rs934197 | 0.901 | 6.9E-09 |
| Esophagus– Muscularis | rs934197 | 0.711 | 1.1E-08 |
| Colon – Sigmoid | rs934197 | 0.730 | 7.3E-07 |
| Muscle – Skeletal | rs56327713 | 0.138 | 8.5E-07 |

